# Cyclic azapeptide cluster of differentiation-36 receptor modulator attenuates left ventricular injury and temporarily reduces long-chain fatty acid accumulation after myocardial ischemia-reperfusion in mice

**DOI:** 10.1101/2025.09.04.674351

**Authors:** Naghme Radmannia, Jade Gauvin, David N. Huynh, Liliane Ménard, Caroline Daneault, Ahsanullah Ahsanullah, André C. Carpentier, William D. Lubell, Matthieu Ruiz, Huy Ong, Sylvie Marleau, Simon-Pierre Gravel

## Abstract

Ischemic heart disease remains a leading global cause of death. We investigated the cardio-protective effects of the selective cluster of differentiation-36 receptor (CD36) modulator azapeptide MPE-298 in a mouse model of myocardial ischemia-reperfusion. Before reperfusion, a single intravenous dose of azapeptide MPE-298 reduced infarct size by 37% and transiently decreased left ventricular (LV) long-chain fatty acid (LCFA) accumulation, independently of saturation status. Metabolomic profiling revealed shifts in amino acids involved in energy production and antioxidant defense. Gene expression analysis showed transient modulation of oxidative stress and inflammation in both heart and adipose tissue. Modulation of CD36 by azapeptide MPE-298 exhibited therapeutic potential for treating acute myocardial ischemia and reperfusion by supporting metabolic recovery and limiting excess LCFA uptake.

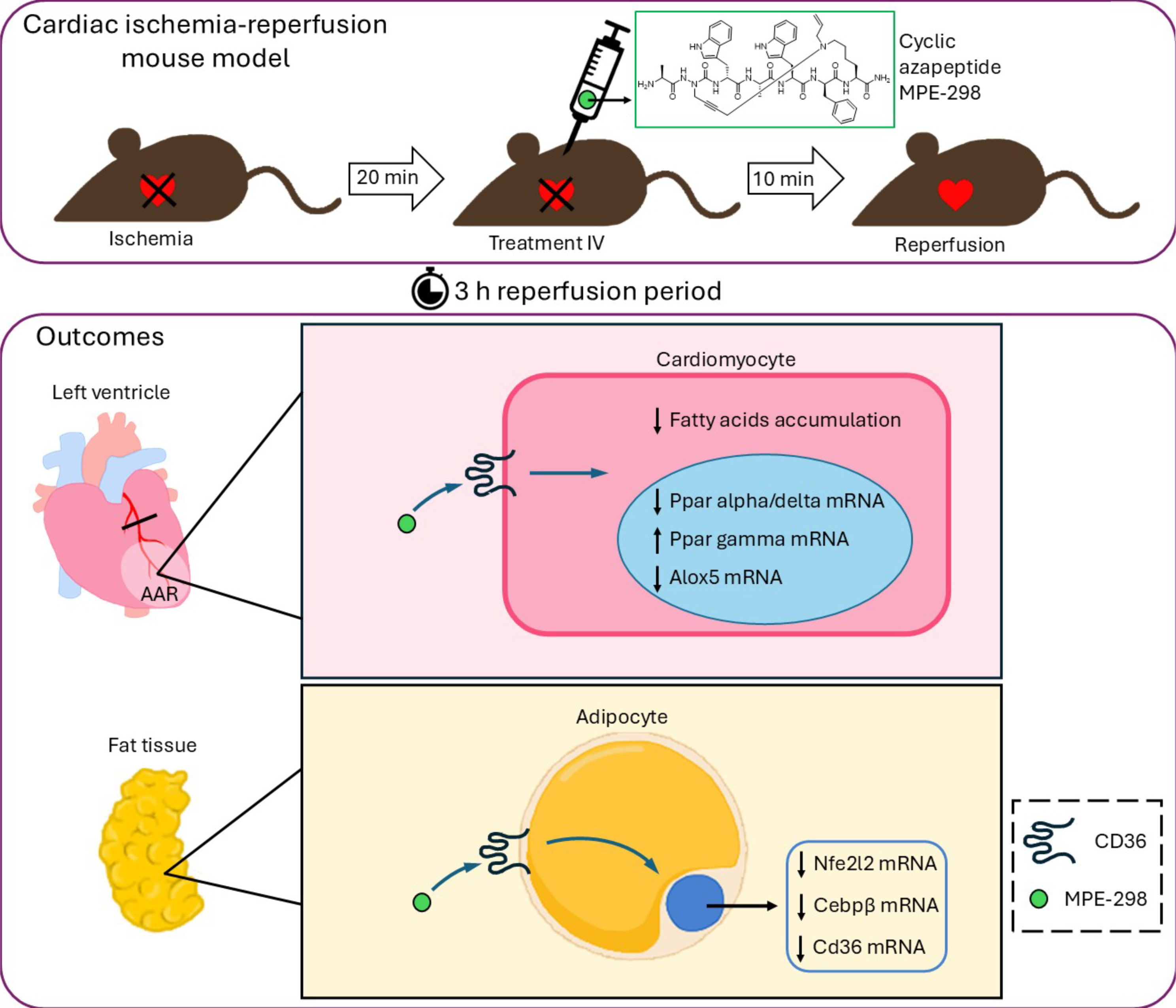

## Introduction

The incidence of cardiovascular diseases, particularly ischemic heart disease (IHD) and stroke, continues to rise globally despite progress in prevention, treatment and monitoring [1]. Deaths related to IHD are expected to persist, largely due to aging populations and the increasing prevalence of hypertension, diabetes, and obesity [2–4] despite reported declines in certain high-income countries [5]. The alarming rise in younger and middle-aged individuals suffering IHD and myocardial infarction (MI) is associated with significant morbidity, psychological effects, and financial strain on patient and family [6]. Among those surviving MI, up to 30% will develop heart failure, despite implementation of pharmacological interventions [7]. Novel therapeutic approaches to treat cardiovascular diseases are urgently needed to remedy the limited long-term effectiveness of current strategies, particularly in the context of prevalent comorbidities and an aging population.

Timely clinical reperfusion is crucial for preserving myocardial tissue and preventing irreversible damage following ischemia. Reperfusion, however, may aggravate tissue injury, due in part to pronounced oxidative and inflammatory consequences [8]. Metabolic disturbances following reperfusion, particularly elevated levels of circulating non-esterified fatty acids (NEFA), can impair recovery of cardiac function [9].

The cluster of differentiation 36 receptor (CD36) is a class B2 scavenger receptor expressed on a range of immune and non-immune cells, where it mediates different cell type-specific functions [10]. In the heart, CD36 facilitates the uptake of free long-chain fatty acids (LCFA) across the plasma membranes of cardiomyocytes [11]. Growth hormone-releasing peptides (GHRPs) have been identified as CD36 ligands [12] and investigated for their potential to protect atherosclerosis [13] as well as for cardioprotective effects in murine models of myocardial ischemia and reperfusion (MI/R) [14]. In models related to myocardial metabolism, the study of GHRPs, such as hexarelin (H-His-D-2-Methyl-Trp-Ala-Trp-D-Phe-Lys-NH_2_), as CD36 ligands has however been limited due to unselective binding to the ghrelin receptor [15]. Selectivity for CD36 has been demonstrated by azapeptide derivatives of GHRP-6 [16, 17].

Cyclic azapeptides, such as MPE-298 have exhibited exceptional CD36 binding affinity and potency in reducing pro-inflammatory nitric oxide and downstream cytokine and chemokine production in macrophages stimulated with a Toll-like receptor-2 agonist [18]. In macrophages, azapeptide MPE-298 was shown to bind and induce CD36 endocytosis through activation of the Lyn and Syk (spleen) tyrosine kinases [19].The internalized CD36–MPE-298 complex inhibited signaling by way of lectin-like oxLDL receptor-1 (LOX-1) activation from oxidized low-density lipoprotein (oxLDL) blocking chemokine ligand 2 (CCL2) secretion, as well as mitochondrial membrane potential depolarization and production of reactive oxygen species (ROS).

Protective effects of azapeptide MPE-298 against atherosclerosis were observed in apolipoprotein E-deficient (apoE^−/−^) mice fed a high-fat high-cholesterol diet [20]. In this study, C57BL/6 wild type mice were administered MPE-298 prior to transient left coronary artery ligation (LCAL) and effects were examined on infarct size and heart and adipose tissue metabolic profile at 3 h and 24 h after reperfusion. Employing lipidomic and metabolomic analysis, changes in energy substrate metabolites were measured in the left ventricular (LV) myocardial tissue.

## Materials and Methods

### Chemicals

Azapeptide MPE-298 was prepared by solid-phase synthesis employing an A^3^-macrocyclization as detailed previously [18]. The cyclic azapeptide demonstrated a strong binding affinity to CD36 (0.1 μM) and exhibited potent activity in reducing the inflammatory response in macrophages.

### Animals

The wild type C57BL/6J inbred mouse strain (stock number 000664) (*Cd36*^+/+^) and CD36-deficient (*Cd36*^-/-^) mice (stock number 019006) were purchased from the Jackson Laboratory (Bar Harbor, ME, US) and bred in specific pathogen-free (SPF) facility. Mice were housed in ventilated cages before being transferred to a controlled environment in static cages. Mice were maintained on a 12 h:12 h light/dark cycle and fed Teklad irradiated global chow (T.2918.15, Inotiv Inc., West Lafayette, IN, US) and water *ad libitum*. All experimental protocols received approval from the Institutional Animal Ethics Committee (No. 23-034) and were conducted in accordance with the guidelines set forth by the Canadian Council on Animal Care and the US National Institutes of Health.

### Experimental groups and protocols

Aged-matched male *Cd36^+/+^* and *Cd36^-/-^* mice (30-35 g) were randomly assigned to one of two main experimental protocols: (i) Both *Cd36^+/+^* and *Cd36^-/-^* mice underwent 30 min of LCAL or sham surgery, followed by intravenous (IV) injection of either 0.9% NaCl (vehicle) or MPE-298, administered 10 min prior to reperfusion. Myocardial infarct area (IA) was assessed 24 h after reperfusion. (ii) For metabolomic and lipidomic analysis, *Cd36^+/+^* mice were subjected to the same LCAL or sham procedures and treatment regimens as in (i) and were euthanized at either 3 h or 24 h following reperfusion.

LCAL was performed following the protocol detailed previously [14] with some modifications. Mice were initially anesthetized with 3.5% isoflurane in 1.5 L/min O_2_, intubated and given 6 mg/kg intraperitoneal lidocaine to prevent arrhythmias. Electrocardiography (ECG) was monitored, and body temperature maintained with a heating pad. Subcutaneous (SC) administration of a 0.9% NaCl - 2.5% dextrose solution (10 mL/kg) was given for the first hour, then 5 mL/kg as needed. Ophthalmic ointment was applied to prevent corneal dryness. Surgery was performed under a Nikon SMZ645 stereomicroscope (0.8 - 5X magnification). A left thoracotomy was performed at the third intercostal space. The transverse and deep pectoral muscles were separated using a retractor to expose the heart. The left coronary artery was visualized under a light block and ligated 1 mm distal to the left atrial appendage using a ½ circle spatula needle and an 8-0 nylon thread was tied over a small piece of Silastic tubing. After ligation, ischemia was maintained during 30 min during which the at-risk region became visibly pale. Ten minutes before reperfusion, mice received an IV injection of 150 μL sterile 0.9% NaCl or 3 μmol/kg MPE-298. After ligation release, the surgical site was sutured, and long-acting buprenorphine (1 mg/kg) was administered subcutaneously followed by extubation. At 3 h or 24 h after reperfusion, mice were euthanized using isoflurane and exsanguinated. Sham-operated mice underwent the same procedure without ligature placement.

### Determination of the myocardial area at risk and infarct size

Twenty-four hours after reperfusion, mice were anesthetized and re-ligated at the original site of LCAL. A 0.5 mL IV injection of 2.0 % Evans blue dye (Sigma-Aldrich, St-Louis, MO, US) was injected retrogradely through the aorta to delineate non-ischemic (blue stained) tissue. Hearts were excised, rinsed with PBS, snap-frozen, and the LV sectioned transversely into 1 mm sections using a mouse heart slicer matrix. Slices were incubated with 1% 2,3,5-triphenyltetrazolium chloride (TTC) at 37°C for 15 min to visualize IA, then fixed in 10% neutral buffered formalin for 12 h. Each slice was weighed and imaged on both sides using a stereomicroscope-mounted digital camera (Coolpix 4500, Nikon). The LV area, area at risk (AAR), and IA were quantified using computerized planimetry with Adobe Photoshop CC 2015, and analysis of treated and untreated groups was performed blindly. Results from each slice were averaged using the weighted mean approach, calculated as (A₁ × W₁) + (A₂ × W₂) + …+ (A_n_ × W_n_), in which A is the IA percentage determined by planimetry, and W is the weight of the corresponding slice [14].

### Targeted FAMEs analysis

The LV were processed for quantitative profiling of bounded fatty acids (FAs) using gas chromatography-mass spectrometry, following established methods [21, 22]. Briefly, approximately 50 mg of pulverized tissue was incubated overnight at 4°C in a 2:1 chloroform/methanol solution containing 0.004% butylated hydroxytoluene (BHT), filtered and dried under nitrogen gas. FAs were analyzed as methyl esters which were prepared by transesterification using acetyl chloride in methanol. The analysis was performed on a 7890B gas chromatograph coupled to a 5977A Mass Selective Detector (Agilent Technologies, Santa Clara, USA), using a J&W Select FAME CP7420 capillary column (100 m x 250 µm inner diameter; Agilent Technologies Inc.) and operated in positive chemical ionization (PCI) mode with ammonia as the reagent gas. Samples were processed under the following conditions: injection at 270 °C in split mode (split ratio: 50:1) and using high-purity helium as the carrier gas at a constant flow rate of 0.44 mL/min. The temperature gradient was set to 190 °C for 25 minutes, followed by an increase of 1.5 °C/min until reaching 236 °C. FAs were analyzed as [M+NH_3_]^+^ ions, and concentrations were determined using standard curves and isotope-labeled internal standards.

### Targeted analysis of amino acids and organic acids

Fifty milligrams of LV tissue were frozen using liquid nitrogen, ground into a powder, and extracted with a solution of 70% methanol and hydroxylamine (1 mol/L) at pH 7.6. The following internal and external isotope-labeled standards were added to the samples: ^13^C_3_-lactic acid (400 nmol), ^13^C_3_-pyruvic acid (20 nmol), ^13^C_4_-β-hydroxybutyric acid (20 nmol), ^13^C_4_-α-ketobutyric acid (1 nmol), D_4_-citrate (20 nmol), ^13^C_4_-α-ketoglutaric acid (20 nmol), D_4_-succinic acid (20 nmol), D_3_-malic acid (40 nmol), ^13^C_4_-acetoacetic acid (20 nmol), ^13^C_4_-fumaric acid (20 nmol), ^13^C_3_-alanine (150 nmol), ^13^C_2_-glycine (50 nmol), ^13^C_5_-valine (20 nmol), ^13^C_6_,^15^N-leucine (10 nmol), ^13^C_6_-isoleucine (10 nmol), ^13^C_5_,^15^N-proline (5 nmol), ^13^C_5_-methionine (15 nmol), ^13^C_5_,^15^N-serine (50 nmol), ^13^C_4_,^15^N-threonine (50 nmol), D_5_-phenylalanine (15 nmol), ^13^C_4_,^15^N-aspartic acid (100 nmol), ^13^C_5_,^15^N-glutamic acid (400 nmol), ^13^C_6_-arginine (150 nmol), ^13^C_9_-tyrosine (10 nmol), ^13^C_6_-histidine (20 nmol), ^13^C_5_,^15^N_2_-glutamine (400 nmol), ^13^C_4_,^15^N_2_-asparagine (15 nmol), ^13^C_11_,^15^N_2_-tryptophan (5 nmol), ^13^C_3_,^15^N-cysteine (40 nmol), and ^13^C_6_,^15^N_2_-lysine (50 nmol). After agitation for 2 min in a sonication bath, the mixture was supplemented with zirconium oxide beads (2.8 mm diameter, Omni International, Kennesaw, GA) and homogenized using Bead Ruptor. Subsequently, the pH of the mixtures was adjusted to between 5 and 6 using hydrochloric acid (1 mol/L). The samples were incubated at 70 °C for 15 min, and centrifuged at 22,000 g for 10 min. The supernatants were evaporated to near dryness. The samples were digested with 100% methanol (2 mL), dried with ammonium sulfate, filtered and rinsed with methanol, a combination of steps repeated twice. After centrifugation at 7,000 g, the supernatants were concentrated and the reduced volumes were transferred into GC-MS vials, and dried. The dry samples were solubilized in pyridine, heated at 45 °C for 90 min, derivatization with *N*-*tert*-butyldimethylsilyl-*N*-methyltrifluoroacetamide at 90 °C for 4 h, and injected into an Agilent 6890N chromatograph coupled with a 5975N mass spectrometer operating in electronic ionization mode using helium as the carrier gas maintained at 7.7 mL/min in split mode (5.6:1). The program temperature was set at 150 °C for 3 min, increased 7 °C/min to 210 °C, held for 3 min, increased 7 °C/min to 310 °C, held for 5.5 min, and finally increased 40 °C/min to 320 °C. Metabolites were identified based on m/z ratios and retention times, and quantified using internal and external standards, and standard curves.

### Gene expression analyses

Total mRNA was extracted from the LV using Trizol^TM^ reagent (Invitrogen Canada Inc., Burlington, Ontario, Canada) followed by solid-phase extraction with Aurum^TM^ total RNA mini kit (Bio-Rad, Mississauga, ON, Canada). Briefly, reverse-transcription to cDNA was performed in a total volume of 20 μL using deoxynucleoside triphosphate (dNTP) nucleotides and Moloney Murine Leukemia Virus (MMLV) Reverse Transcriptase (Invitrogen) at 37°C for 75 min followed by 95°C for 10 min. The sample volumes were diluted 1:10 in diethylpyrocarbonate (DEPC)-treated water to remove RNases and kept at –20°C until use. Realtime quantitative polymerase chain reaction (qPCR) amplification was carried out in a volume of 10 μL with 1 μL of cDNA, 20 μM of each specific primer and SsofastTM EvaGreen® Supermix (BioRad Laboratories, Hercules, CA, USA). The cycling protocol consisted of 40 cycles starting at 95°C for 30 s, then at 60°C for 90 s, and at 95°C for 15 s using a QuantStudio 5 realtime PCR system. Relative mRNA expression levels were determined using the comparative cycle threshold (Ct) method (2^−ΔΔCt^) and normalized to the mean of the reference genes *Ppia* and *Hprt* unless otherwise specified. The primer sequences are detailed in Table 1. Results are presented as fold changes relative to the corresponding sham-treated mice.

**Table 1.**
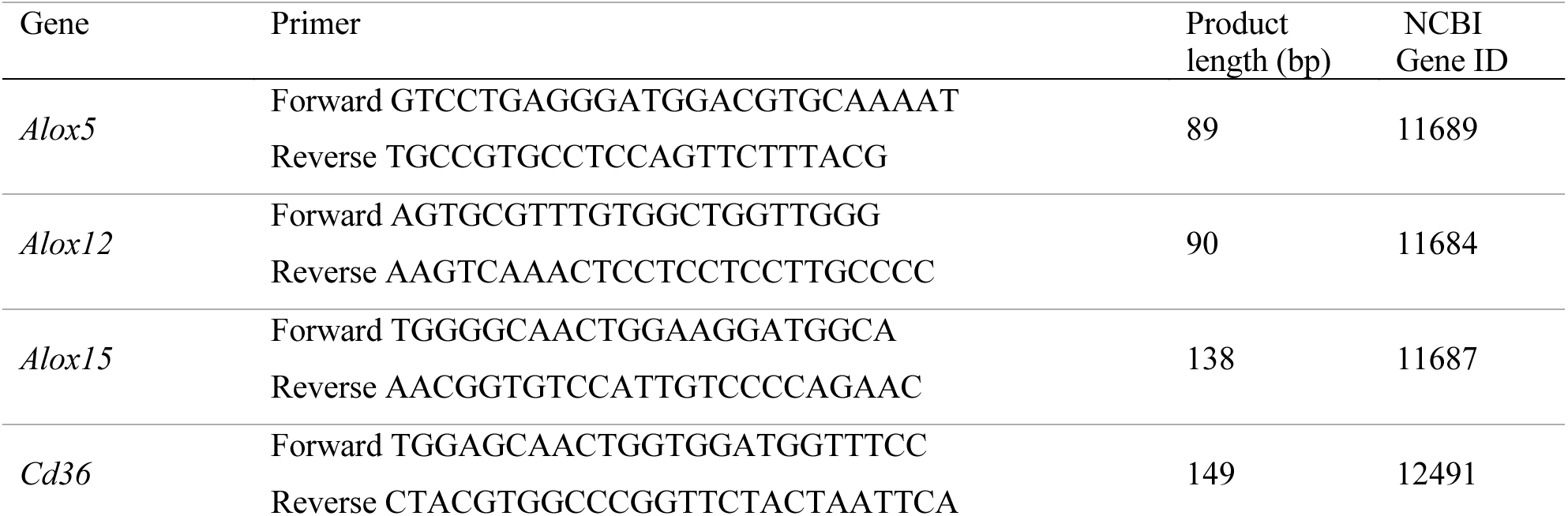
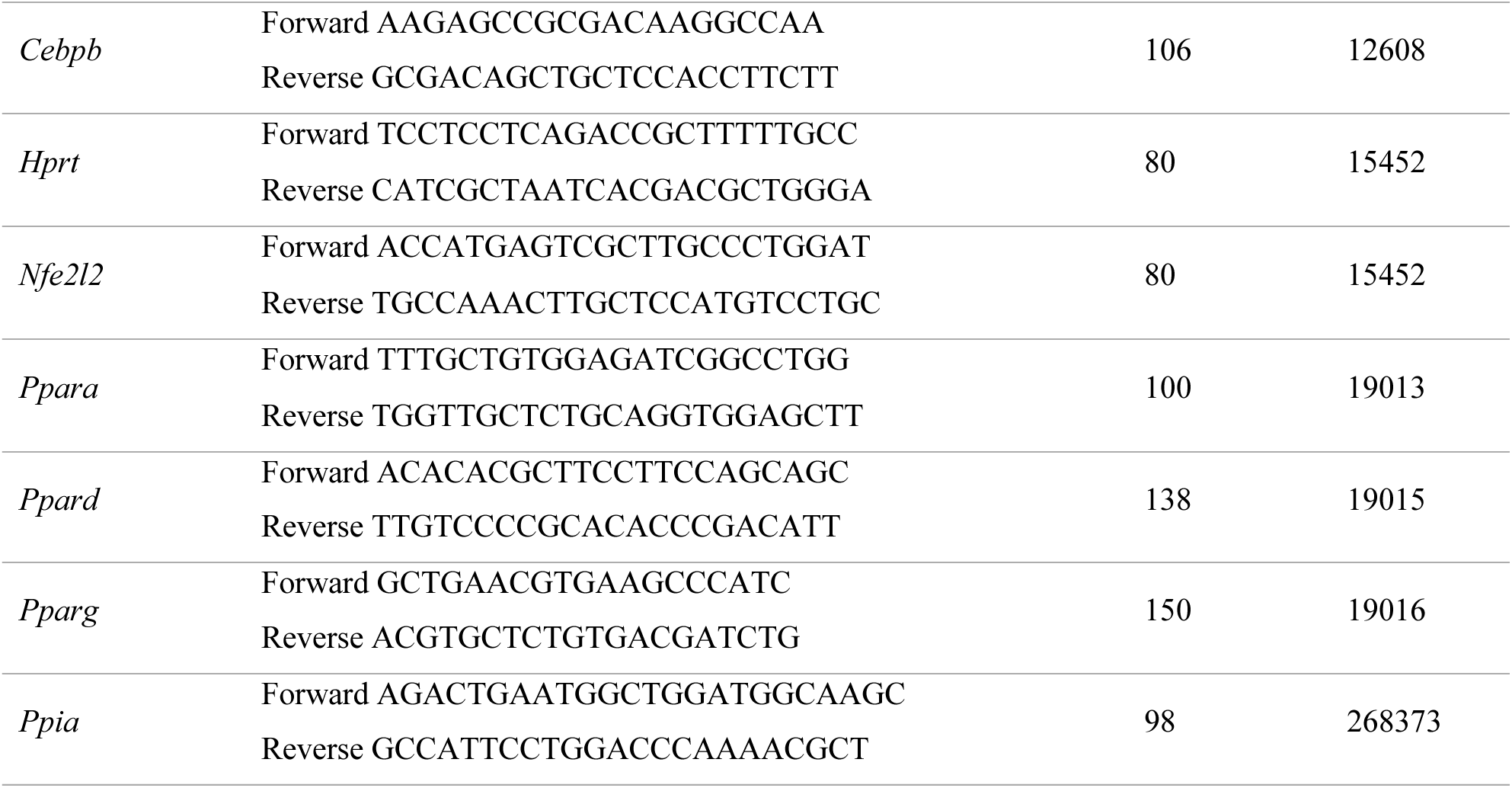
Murine primer sequences for qPCR.

### Statistical analyses

All data were analyzed using GraphPad Prism (version 10.4.2, San Diego, CA, USA) and are presented as mean ± SEM. Statistical comparisons were performed using either Student’s *t*-test or two-way ANOVA where appropriate, with time and treatment as factors, followed by Fisher’s post hoc test for multiple comparisons, as detailed in the figure legends. Normality was assessed using Shapiro-Wilk and Kolmogorov-Smirnov tests prior to conducting *t*-test (GraphPad Prism). Outliers identified by Grubbs’ test (α = 0.05), were excluded, as indicated in the figure legends. Statistical significance was defined as p < 0.05.

## Results and discussion

### Azapeptide MPE-298 reduced left ventricular injury after myocardial ischemia and reperfusion

The cardioprotective effect of azapeptide MPE-298 was initially investigated by administration of a single IV dose shortly before reperfusion. Hypothesizing that azapeptide MPE-298 would act in a CD36-dependent manner, both *Cd36^⁺/⁺^* and *Cd36^⁻/⁻^* mice were subjected to LCAL and reperfusion for 24 h. Ten minutes before reperfusion, mice received an IV injection of either vehicle (0.9% NaCl) or azapeptide MPE-298 (3 μmol/kg) (Fig. 1A). The selected dose was based on prior dose-response experiments, which demonstrated a dose-dependent reduction in infarct size [23].

**Fig 1.**
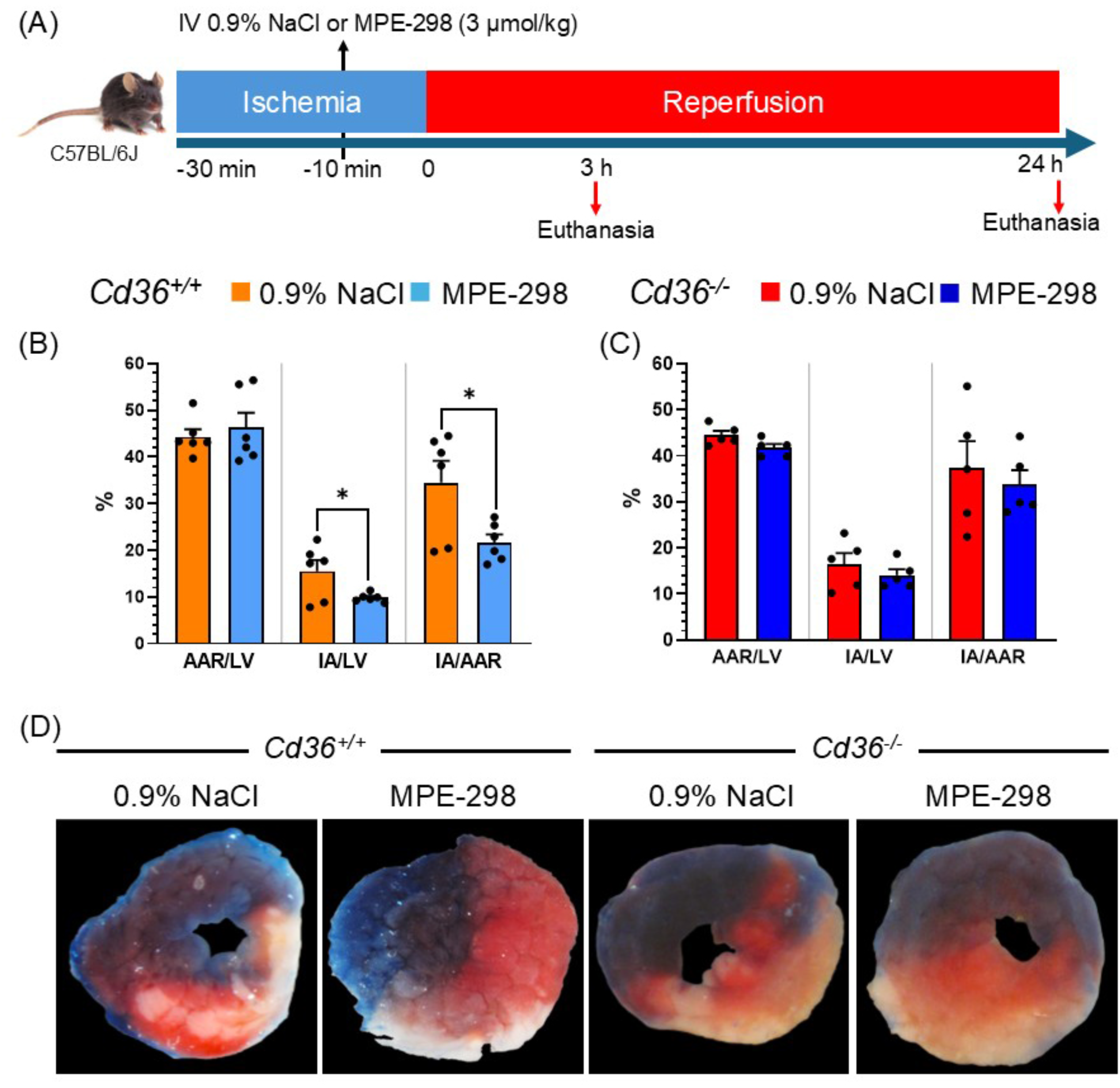
Treatment with MPE-298 reduces infarct area in *Cd36*^+/+^ mice but not in *Cd36^-/-^* mice after MI/R (A) Experimental timeline: mice underwent LCAL for 30 min, followed by reperfusion and monitoring at 3 h and 24 h. Azapeptide MPE-298 (3 μmol/kg) or vehicle (0.9% NaCl) were administered IV 10 min before reperfusion. (B,C) Quantification of myocardial injury at 24 h post-MI/R: bar graphs and dot plots show (i) area at risk (AAR) as a percentage of left ventricular (LV) area, (ii) infarct area (IA) as a percentage of LV area, and (iii) IA as a percentage of AAR in *Cd36^+/+^* (n = 6/group) and *Cd36^-/-^* (n = 5/group) mice. (D) Representative TTC- and Evans blue-stained mid-ventricular cross-sections from *Cd36^+/+^* and *Cd36^-/-^* mice showing the AAR (red), infarcted tissue (white), and non-ischemic tissue (blue) 24 h after reperfusion. Data are mean ± SEM. * p < 0.05, as assessed by unpaired Student’s *t*-test.

The AAR, relative to LV, was consistent across genotypes and treatment groups, averaging 44%, indicating uniformity in the extent of ischemic insult and effective experimental reproducibility (Fig. 1B and 1C). In *Cd36^⁺/⁺^* mice, treatment with MPE-298 significantly reduced IA, as reflected by a 37% decrease in both IA/LV and IA/AAR ratios compared to vehicle-treated mice (p < 0,05; Fig. 1B). In contrast, no significant IA reduction was observed in *Cd36^⁻/⁻^* mice (Fig. 1C), indicating that the cardioprotective effect of MPE-298 is CD36-dependent. The protective effect was further illustrated in TTC-stained mid-ventricular sections in which MPE-298-treated *Cd36^+/+^*mice showed visibly smaller infarct size compared to those from mice treated with vehicle or from *Cd36^⁻/⁻^* mice regardless of treatment (Fig. 1D).

Administration of a single dose of azapeptide MPE-298 shortly before reperfusion provided acute protection against LV injury in a CD36-dependent manner. In related earlier studies, a 14-d pretreatment with a GHRP and selective linear azapeptide CD36 ligands EP 80317 and CP-3(iv) prior to 30 min ischemia was shown to respectively reduce the IA to LV area ratio by 34% and 56% [14, 23]. Such attenuations of LV injury were primarily attributed to a reduced myocardial lipid burden associated with lower plasma levels of circulating NEFA. The CP-3(iv) pretreated mice were also shown to exhibit increased expression of adiponectin in adipose tissue and higher circulating adiponectin levels [14, 23]. Moreover, administration of linear azapeptide CP-3(iv) 10 min before reperfusion caused a dose-dependent reduction of the IA to LV area ratio up to 44% at the highest dose tested (1000 nmol/kg) compared with vehicle-treated CD36^+/+^ mice.

### Azapeptide MPE-298 attenuates LCFA accumulation in LV after myocardial ischemia and reperfusion

Cellular uptake of LCFA is primarily governed by CD36 at the muscle fiber cell membrane, the sarcolemma [24]. The effects of azapeptide MPE-298 on FA composition and abundance were consequently examined in the LV after reperfusion. Following MI, reperfusion of salvageable ischemic myocardium leads to a rapid restoration of FA uptake and oxidation [9]. This early rise in LCFA β-oxidation disrupts the balance between glucose and FA oxidation as competing sources of adenosine triphosphate (ATP) production [25]. The predominance of β-oxidation requires more oxygen for ATP production, and leads to excess hydrogen ions production, which in turn contributes to ionic imbalances that decrease heart efficiency [26]. Transient inhibition of early uptake of LCFA after reperfusion may protect the heart from detrimental oxidative stress after reperfusion and improve contractile recovery. Consistent with this hypothesis, mice pretreated with EP 80317 were shown to exhibit reduced myocardial FA uptake 6 h after reperfusion by micro-positron emission tomography (µPET) using [^18^F]-labelled fluoro-6-thia-heptadecanoic acid as a tracer of NEFA [14].

Targeted analysis of FAs was performed after esterification as methyl esters (FAMEs). Analysis of fold changes in the LV revealed that, at 3 h post-reperfusion, mice receiving a single IV dose of the azapeptide MPE-298 showed transient reductions in certain LCFAs compared with vehicle-treated controls. The decrease was observed across all classes of saturation of FAs: saturated (SFA), monounsaturated (MUFA), and polyunsaturated (PUFA, heatmap Fig. 2A). Significant reductions were observed compared to those of vehicle-treated mice in the relative abundance of SFA [myristic, pentadecanoic and palmitic acids (Fig. 2B-D)], MUFA [palmitoleic and vaccenic acids (Fig. 2E and 2F)], and PUFA [arachidonic, eicosatrienoic (ETE) and docosahexaenoic (DHA) acids (Fig. 2H-J)]. A comparable downward trend was noted for oleic acid (Fig. 2G), and modest downward trends were observed for docosapentaenoic (DPA) and linoleic acids (Fig. 2K and 2L). SFAs have long been considered detrimental to cardiovascular health, and dietary recommendations to limit their intake have persisted. Although higher total SFA concentrations were associated with increased cardiovascular risk, further analysis revealed positive or negative associations according to SFA subtypes [27]. Notwithstanding, the long-term health effects of SFA and certain MUFA remain a subject of ongoing debate, whereas the benefits of PUFA are more widely accepted [28], albeit with some conflicting evidence [29]. Importantly, the changes in LCFA LV content following treatment with azapeptide MPE-298 were temporary, and most FA were normalized by 24 h post-reperfusion. At this time point, levels of vaccenic acid and DPA were higher than those in vehicle-treated mice. Notably, levels of SFA, MUFA, and linoleic acid declined between 3 h and 24 h in vehicle-treated mice to sham values, suggesting that the peak in LCFA uptake occurs primarily during the early reperfusion phase, consistent with previous reports [30]. In sum, azapeptide MPE-298 reduced transiently myocardial LCFA content independent of saturation state in the LV of mice subject to myocardial infarction and reperfusion (MI/R). Previous findings in mice pretreated with the CD36 ligand EP80317, showed transient suppression of peripheral lipolysis and reduced circulating NEFA levels at 6 hours following MI/R [14]. In the present study, plasma NEFA levels showed a modest but non-significant decline at 3 hours, while plasma TG remained unchanged. Consistent with our earlier findings linking reduced lipolysis to lower cardiac FA burden, this prompted us to assess markers of lipolysis in epididymal fat.

**Fig. 2.**
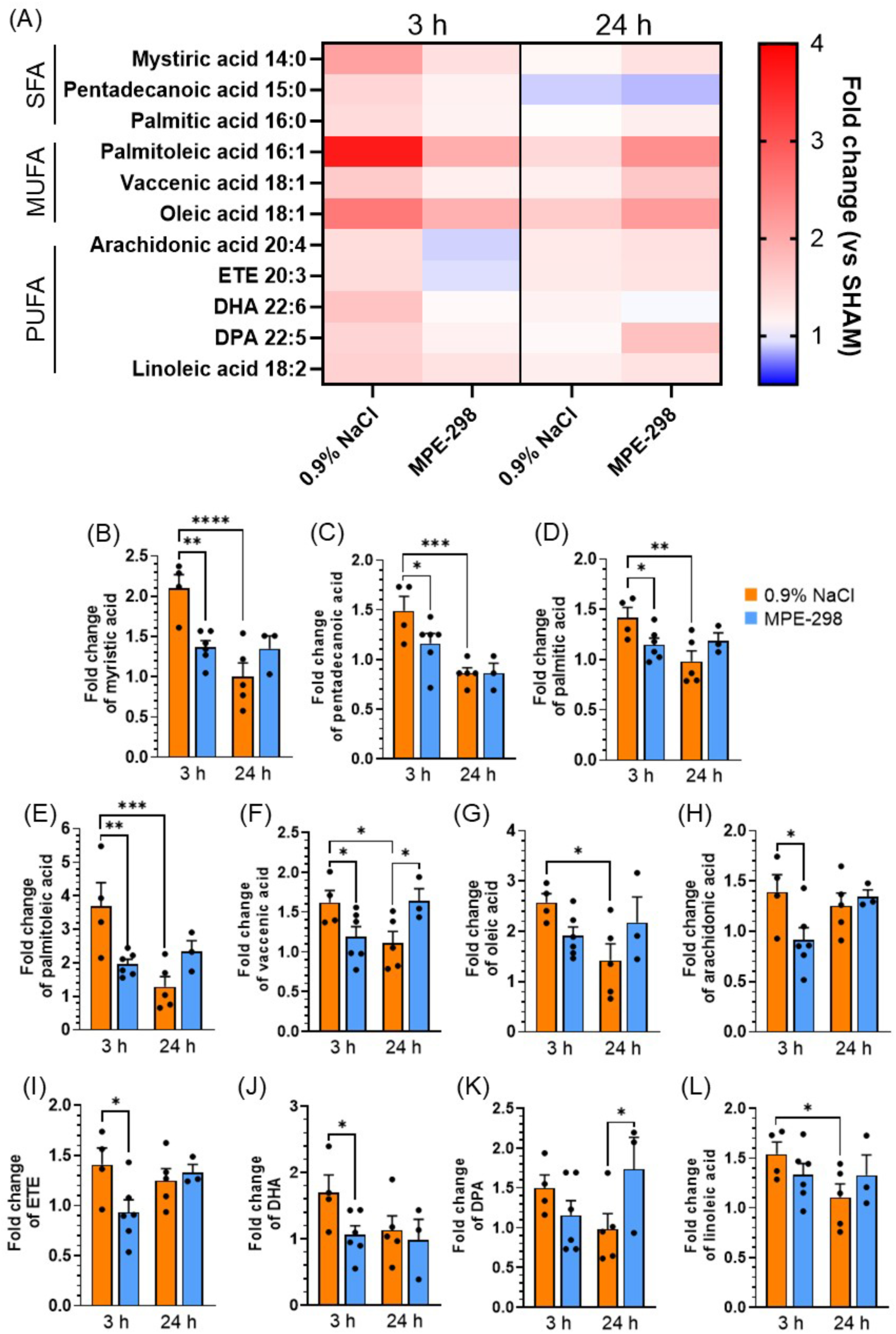
Administration of a single IV dose of azapeptide MPE-298 (3 μmol/kg) prior to reperfusion induces a transient reduction of the relative left ventricular (LV) content of LCFA at 3 h post-reperfusion in mice that underwent a temporary LCAL for 30 min. (A) Heatmap depicting the relative LCFA content in the LV of vehicle- and MPE-298-treated mice at 3 h and 24 h post-reperfusion, with data normalized to corresponding sham-operated controls. Bar graphs represent the relative LV content of specific LCFAs including (B) myristic acid, (C) pentadecanoic acid, (D) palmitic acid, (E) palmitoleic acid, (F) vaccenic acid, (G) oleic acid, (H) arachidonic acid, (I) eicosatrienoic acid (ETE), (J) docosahexaenoic acid (DHA), (K) docosapentaenoic acid (DPA) and (L) linoleic acid. Data are expressed as the mean ± SEM of 4-6 mice. Statistical analysis was performed using a 2-way ANOVA with treatment and time as independent variables, followed by Fisher’s post-hoc test. * p < 0.05.

### Azapeptide MPE-298 alters expression of lipolysis genes in epidydimal adipose tissue post-myocardial ischemia and reperfusion

Lipolysis is acutely activated in response to MI/R-induced oxidative stress and inflammation [31]. The transcription factor nuclear factor erythroid-derived 2-like 2 (Nfe2l2) is a key regulator of cellular antioxidant defense mechanisms which are activated in response to cellular stress, such as those during MI/R injury [32]. Early activation of Nfe2l2 has however been shown to exacerbate MI/R injury by promoting myocardial inflammation, release of pro-inflammatory cytokines and myocardial macrophage inflammatory phenotypes [33]. Mice that were administered a single IV dose of azapeptide MPE-298 (3 μmol/kg) exhibited accordingly 3 h post-reperfusion a transient reduction in *Nfe2l2* gene expression in epididymal adipose (Fig. 3A) accompanied by a reduction of the downstream target genes for the transcription factor C/EBPβ (Fig 3B) and for CD36 (Fig. 3C). The transcription factor C/EBPβ has been shown to transcriptionally upregulate CD36 mRNA expression [34]. Reduced CD36 expression has been linked to decreased lipolytic breakdown of lipids by lipases [35]. The functional relevance of the expression of CD36 in adipose tissue is supported by the reduction of lipolysis on treatment with the CD36 inhibitor sulfo-N-succinimidyl oleate [35]. Notably, expression levels of Nfe2l2 and its targets genes recovered fully at 24 h after reperfusion (Fig. 3A–C). In contrast, a 14-day pre-treatment with the linear azapeptide CP-3(iv) at a lower dose (300 nmol/kg), led to elevated expression of Nfe2l2 and its downstream targets, including CCAAT/enhancer-binding protein beta (Cebpb) and Cd36 in epidydimal fat at 6 h post-reperfusion [23]. These discrepancies may be attributed to differences in treatment, treatment duration, sampling time, and administered dose.

**Fig. 3.**
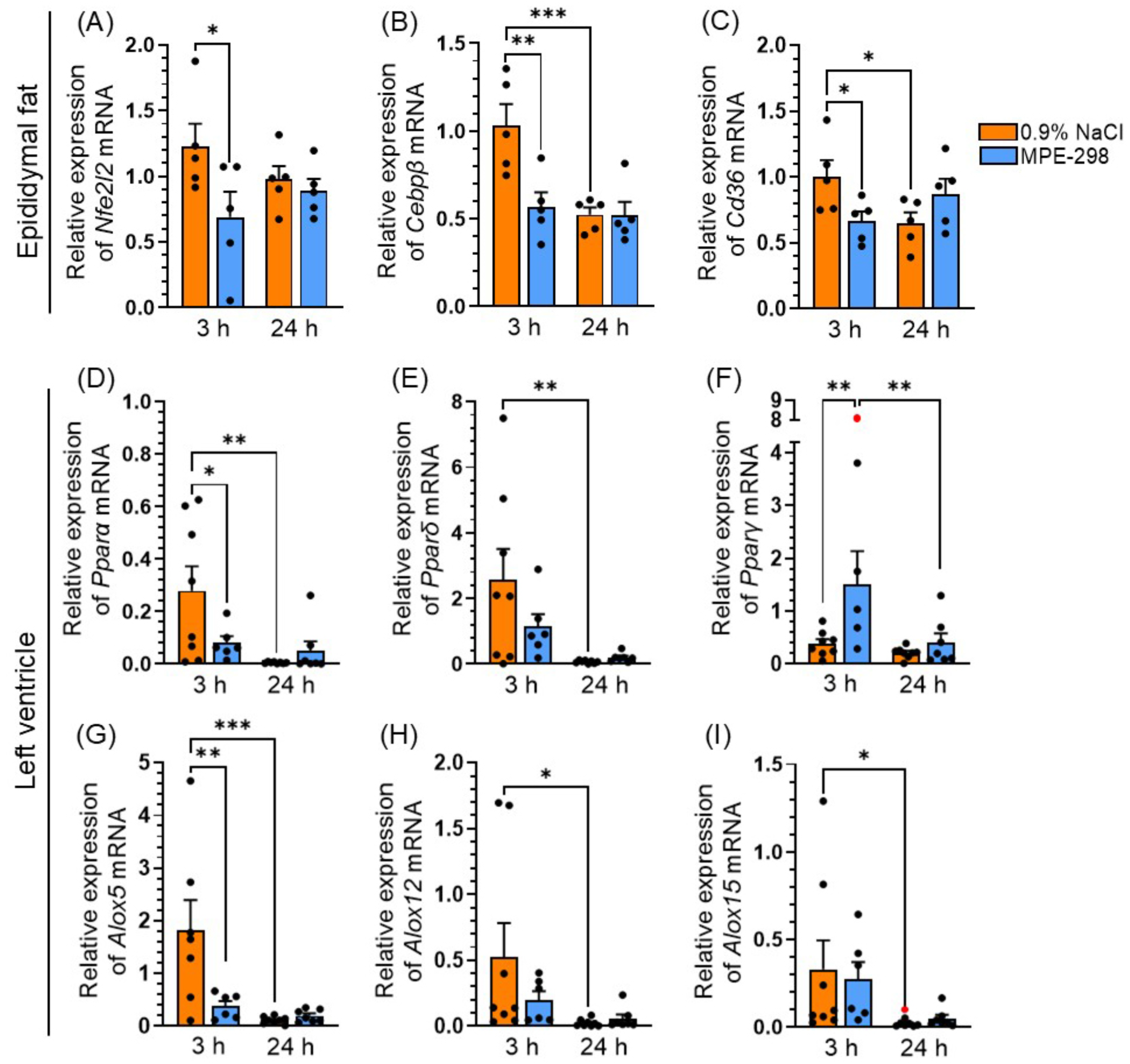
Administration of azapeptide MPE-298 reduces the expression of genes involved in metabolic and inflammatory pathways. Mice were treated IV 10 min prior to reperfusion with either 0.9% NaCl or MPE-298 (3 μmol/kg) and monitored at 3 h or 24 h post-reperfusion. Levels in adipose tissue of mRNA: (A) *Nfe2l2*, (B) *Cebpβ* and (C) *Cd36*, and in left ventricle (LV): (D) *Pparα*, (E) *Pparδ*, (F) *Pparγ*, (G) *Alox5*, (H) *Alox12* and (I) *Alox15*. Data are expressed as the mean ± SEM of 5-8 mice. Statistical analysis was performed using a 2-way ANOVA with treatment and time as independent variables, followed by Fisher’s post-hoc test. Two outliers (red dots) were identified using the Grubbs’ test. * p < 0.05.

The LCFA serve as endogenous ligands of peroxisome proliferator-activated receptor (PPAR) subclasses, with PPARα being the most abundantly expressed isoform in the heart [36]. Playing a key role in regulating mitochondrial lipid β-oxidation, PPARα promotes fatty acid oxidation (FAO) and concurrently reduces glucose utilization in cardiomyocytes [36]. Associated with inflammation and elevated myocardial ROS, MI/R leads to upregulation of PPARα expression in the myocardium. Temporary reduction of PPARα expression may be beneficial during the early hours of reperfusion because enhancement of β-oxidation may impair cardiac efficiency. Previously, pretreatment with linear azapeptide CP-3(iv) reduced myocardial ROS levels in mouse LV homogenates [23]. Accordingly, mice treated with cyclic azapeptide MPE-298 exhibited reduced PPARα gene expression at 3 h post-reperfusion after LCAL, compared to vehicle-treated animals (Fig. 3D). Similar to PPARα, PPARδ has been implicated in promoting myocardial FAO [37]. At 3 h after reperfusion, the LV of mice treated with azapeptide MPE-298 showed a trend towards decreased PPARδ gene expression (Fig. 3E), but transiently elevated PPARγ mRNA expression (Fig. 3F). Although associated with the upregulation of genes involved in FAO during MI/R injury, PPARγ is primarily expressed in adipose tissue and macrophages [38]. After reperfusion, antioxidant and anti-inflammatory responses may become predominant [38, 39], supporting a role for PPARγ as a key transcription factor mediating anti-inflammatory responses following MI/R.

Animal models have controversially shown that myocardial MI/R results in elevated production of leukotrienes in the myocardium [40]. Growing evidence indicates that metabolites derived from the 5-lipoxygenase (Alox5) pathway contribute negatively to myocardial functional recovery. Previously, pretreatment with EP 80317 was found to selectively downregulated Alox5 gene expression and to reduce leukotriene B_4_ (LTB_4_) levels in the lungs of mice subjected to transient skeletal ischemia-reperfusion, suggesting potential for modulating the inflammatory response [41]. Noting the transient reduction of levels of the LCFA arachidonic acid in the LV at 3 h after reperfusion (Fig. 2H), we investigated the effect of azapeptide MPE-298 on the Alox5 mRNA levels following MI/R. Azapeptide administration led to a transient reduction in Alox5 mRNA expression in mice LV (Fig. 3G), without significantly changing transcript levels of Alox12 (Fig. 3H) and Alox15 (Fig. 3I). In the early response to MI/R, metabolites from Alox5 oxidation may play roles which are transiently suppressed upon azapeptide treatment. Previously, LTB_4_, a lipid mediator of inflammation was shown to be an endogenous ligand for PPARα [42]. Reduction in Alox5 expression at 3 h after reperfusion may decrease transiently activation of PPARα. The mechanistic links between ALOX5, CD36 and PPARα warrant further investigation in the context of MI/R injury.

### Azapeptide MPE-298 alters amino acid composition in the left ventricle following myocardial ischemia and reperfusion

In the early phase of reperfusion following ischemia, we observed that the LV of mice treated with MPE-298 exhibited distinctly reduced levels of LCFA (Fig. 2). Previously, energy metabolism in the heart after reperfusion was shown to shift towards a greater reliance on glucose [30] and potentially anaplerosis of specific amino acids [43]. In addition to serving as metabolic substrates in MI/R injury, certain amino acids may play broader roles, such as modulating the cellular responses to stress. The influence of azapeptide MPE-298 on levels of amino acids and tricarboxylic acid (TCA) cycle intermediates was examined in the LV of mice subjected to MI/R (heatmaps Fig. 4A and 4B). Among TCA metabolites, plasma levels of 2-hydroxyglutarate, fumarate and malate have been associated with increased cardiovascular risk [44], while the relative levels of these intermediates in the LV tended to decrease in MPE-298-treated mice (Fig. 4B). Amino acid glutamate serves as a substrate that fuels the TCA cycle through conversion to α-ketoglutarate via transamination (Fig. 4C) [45]. Glutamine (Fig. 4D) and proline (Fig. 4E), which can be metabolized to glutamate [46], have been similarly shown to contribute to the production of α-ketoglutarate [47]. Glutamate, cysteine and glycine are components in the synthesis of glutathione, a potent intracellular antioxidant [48]. In the LV of mice treated with azapeptide MPE-298 at 3 h after reperfusion, glutamate and cysteine declined transiently, and glutamine and proline both showed modest trends toward reduction (Fig. 4E and 4F). Moreover, the aromatic amino acids, tyrosine (Fig. 4H) and phenylalanine (Fig. 4I) were reduced at 3 h and 24 h in MPE-298-treated compared to vehicle-treated mice. Although metabolism of tyrosine and phenylalanine by way of aromatic ring hydroxylation may lead to the TCA intermediate fumarate [49], levels of the latter were not significantly altered (Fig. 4B). Amino acids serve many roles beyond energy metabolism, including protein synthesis, cofactors in biochemical reactions and signaling intermediates [50]. For example, a trend toward decreased LV levels was observed for alanine (Fig. 4J), which undergoes transamination to pyruvate [43] but neither pyruvate (Fig. 4K) nor lactate (Fig. 4L) exhibited significant differences in MPE-298-treated and vehicle-treated mice suggesting little perturbation of this metabolic pathway early after reperfusion.

**Fig. 4.**
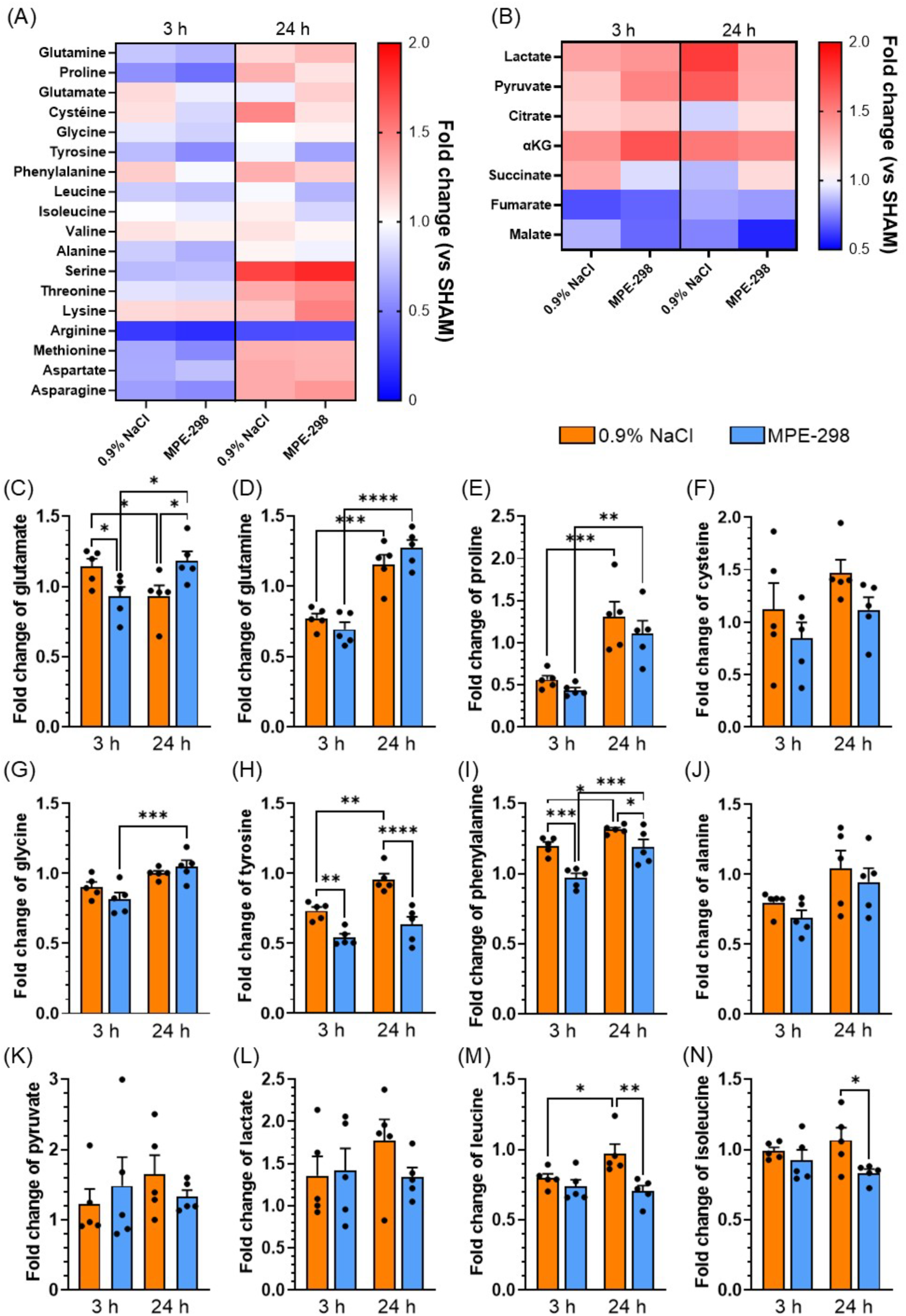
A single IV dose of azapeptide MPE-298 (3 μmol/kg) administered before reperfusion induced a transient reduction in the left ventricular (LV) content of specific amino acids at 3 h post-reperfusion in mice that underwent a temporary LCAL for 30 min. Heatmaps depicting fold change relative to corresponding sham-operated mice of (A) amino acids and (B) TCA intermediates in the LV of vehicle- and MPE-298-treated mice at 3 h and 24 h post-reperfusion. Bar graphs represent the relative LV content of specific amino acids and metabolites: (C) glutamate, (D) glutamine, (E) proline, (F) cysteine (G) glycine (H) tyrosine, (I) phenylalanine, (J) alanine, (K) pyruvate, (L) lactate, (M) leucine and (N) isoleucine. Data are expressed as the mean ± SEM of 4-5 mice. Statistical analysis was performed using a 2-way ANOVA with treatment and time as independent variables, followed by Fisher’s post-hoc test. * p < 0.05.

The branched-chain amino acids (BCAA) leucine, isoleucine, and valine are essential and must be obtained through diet. They contribute minimally to normal cardiac energy production, but MI/R impairs their catabolism leading to BCAA accumulation, perturbed cardiac metabolic homeostasis and stress [51, 52]. Although no significant changes in LV content of BCAA were observed between vehicle and MPE-298 groups at 3 h after reperfusion (Fig. 4M-N), by 24 h, both leucine and isoleucine levels were reduced in the latter, suggesting little accumulation in the myocardium but potential roles in myocardial metabolism in the critical early period after reperfusion.

The limitations of this study are due in part to examination of only two time points after reperfusion and only male animals. Despite these constraints, cardioprotective effects were evident after administration of a single dose of azapeptide MPE-298 prior to reperfusion. The study revealed the early beneficial impact of azapeptide treatment on myocardial metabolism specifically the reduction of LCFA following MI/R. The beneficial effects on myocardial metabolism complement earlier findings from studies featuring pretreatment with a CD36 ligand which respectively reduced and increased circulating NEFA and adiponectin levels with increased expression of the latter in adipose tissue [14, 23]. Selective CD36 targeting has shown additional therapeutic benefit for treating MI/R injury.

In conclusion, azapeptide MPE-298 exhibited beneficial effects after myocardial reperfusion of ischemic hearts in a CD36 dependent manner. Metabolic analysis indicated that MPE-298 caused a transient reduction of LCFA levels in the LV early after reperfusion of treated mice. This reduction was associated with downregulation of key metabolic and inflammatory pathways, including the β-oxidation pathways regulated by PPARs. For example, diminished arachidonic acid levels in the LV correlated with a transient reduction in Alox5 mRNA expression. Moreover, a transient reduction in certain amino acids was also observed at 3 h after reperfusion. By 24 h post-reperfusion, the levels of LCFA and most amino acids in the LV returned to values comparable to vehicle-treated controls, with the notable exception of BCAA which remained relatively lower. Collectively, these results support the hypothesis that a CD36 ligand may exert cardioprotective effects by modulating both metabolic and antioxidant anti-inflammatory pathways.

## Acknowledgments

This work was supported by the Canadian Institutes of Health Research (PJT - 178227), the Heart and Stroke Foundation of Canada (G-18-0022167), an educational grant from Mperia Therapeutics Inc., Natural Sciences and Engineering Research Council of Canada Discovery Grants (#04079, #04423, and #06647), and the Fonds de Recherche du Québec - Nature et Technologies from the Centre in Green Chemistry and Catalysis (FRQNT-2020-RS4-265155-CCVC). JG is a recipient of a scholarship from the Fonds de recherche du Québec – Santé (FRQS).

## Author contributions

SM conceived and designed experiments; SPG, SM and HO supervised the studies; WDL and AA provided resources; NR, JG, LM and DNH performed experiments and analyzed the data; CD and MR performed GC-MS and provided results. SPG, SM and WDL wrote the manuscript; WDL edited the manuscript and ACC, HO, DNH, LM, NR and JG made manuscript revisions.

## Abbreviations

AAR: area at risk
ALOX5: arachidonate 5-lipoxygenase
CCL2: chemokine ligand 2
CD36: cluster of differentiation 36
Cebpβ: CCAAT/enhancer-binding protein beta
Ct: cycle threshold
DEPC: diethylpyrocarbonate
DHA: docosahexaenoic acid
dNTP: deoxynucleoside triphosphate
DPA: docosapentaenoic
ETE: eicosatrienoic acid
FA: fatty acid
IA: infarct area
IHD: ischemic heart disease
GHRPs: growth hormone-releasing peptides
IV: intravenous or intravenously
LCAL: left coronary artery ligation
LCFA: long-chain fatty acids
LOX-1: lectin-like oxidized low-density lipoprotein receptor-1
LTB_4_: leukotriene B_4_
LV: left ventricle or left ventricular
MI: myocardial infarction
MI/R: myocardial ischemia and reperfusion
MMLV: Moloney murine leukemia virus
MUFA: monounsaturated fatty acid
NEFA: non-esterified fatty acids
Nfe2l2: nuclear factor erythroid-derived 2-like 2
oxLDL: oxidized low-density lipoprotein
PCI: positive chemical ionization
PPAR: peroxisome proliferator-activated receptor
PUFA: polyunsaturated fatty acid
qPCR: quantitative polymerase chain reaction
ROS: reactive oxygen species
SFA: saturated fatty acid
SC: subcutaneous
TCA: tricarboxylic acid
TTC: 2,3,5-triphenyltetrazolium chloride
µPET: micro-positron emission tomography

